## Supplemental Data for "Understanding Painful versus Non-Painful Dental Pain in Male and Female Patients: A Transcriptomic Analysis of Human Biopsies"

| **Genes Upregulated in Symptomatic Males Compared to Asymptomatic Males** | |
| --- | --- |
| **Genes** | **Function** |
| VNN1 | Immune Response |
| IL10 | Immune Response |
| GNLY | Immune Response |
| PLK2 | Multiple Functions |
| GUCY1A3 | Other |
| TDO2 | Other |

**Table 1**

| **Genes Downregulated in Symptomatic Males Compared to Asymptomatic Males** | |
| --- | --- |
| **Genes** | **Function** |
| TWIST2 | Immune Response |
| SAA2 | Immune Response |
| TIMP3 | Immune Response |
| CXCL14 | Immune Response |
| CCL22 | Immune Response |
| SOD3 | Repair |
| IGF2 | Repair/Regeneration |
| ITGBL1 | Cell Adhesion |
| FBLN5 | Vascular and Cell Adhesion |
| COL8A1 | Extracellular Matrix and Vascular |
| CORIN | Neural |
| LEPR | Neural and Bone Metabolism |
| MOXD1 | Other |
| OMD | Other |
| DIRAS3 | Other |
| CPXM2 | Other |
| LTBP2 | Other |
| DKK3 | Other |

**Table 2**

| **Genes Upregulated in Symptomatic Females Compared to Asymptomatic Females** | |
| --- | --- |
| **Genes** | **Function** |
| PVRIG | Immune response |
| CCR7 | Immune Response |
| ERAP2 | Immune Response |
| WFDC2 | Multiple Functions |
| DCLK1 | Neural |
| F3 | Vascular |
| LAMA2 | Extracellular Matrix |
| HERC2P2 | Other |
| LOC100130231 | Other |
| LOC144571 | Other |
| SPOCK2 | Other |

**Table 3**

| **Genes Downregulated in Symptomatic Females Compared to Asymptomatic Females** | |
| --- | --- |
| **Genes** | **Function** |
| STAP1 | Immune Response |
| DAPK2 | Immune Response |
| MMP13 | Extracellular Matrix |
| SNORA53 | Other |
| SNORD33 | Other |
| SQLE | Other |
| HIST1H3B | Other |
| IBSP | Other |
| SNORD110 | Other |
| RNU12 | Other |
| HIST1H3C | Other |
| HIST1H2BG | Other |
| ADCYAP1 | Other |
| C3orf70 | Other |

**Table 4**

| **Genes Upregulated in Symptomatic Females Compared to Symptomatic Males** | |
| --- | --- |
| **Genes** | **Function** |
| ERAP2 | Immune Response |
| CHIT1 | Immune Response |
| C3 | Immune Response |
| SIGLEC1 | Immune Response |
| SLC18A2 | Neural |
| CADM3 | Cell Adhesion |
| CTGF | Cell Adhesion |
| LRRC15 | Cell adhesion |
| ABI3BP | Extracellular Matrix |
| CHRDL2 | Bone Metabolism |
| SOD3 | Repair |
| SIK1 | Other |
| CPZ | Other |
| FOLR2 | Other |
| ACAB | Other |

**Table 5**

| **Genes Downregulated in Symptomatic Females Compared to Symptomatic Males** | |
| --- | --- |
| **Genes** | **Function** |
| PDPN | Immune Response |
| VNN1 | Immune Response |
| IL1B | Immune Response |
| MIR650 | Multiple Functions |
| LRRK2 | Neural |
| HIST1H3J | Other |
| HIST1H2AB | Other |
| PGM2 | Other |
| PGD | Other |
| CYorf15A | Other |
| CYorf15B | Other |

**Table 6**

| **Genes Upregulated in Asymptomatic Females Compared to Asymptomatic Males** | |
| --- | --- |
| **Genes** | **Function** |
| HLA-J | Immune Response |
| CX3CR1 | Immune Response |
| ATP10B | Neural |
| COCH | Extracellular Matrix |
| RNU12 | Other |
| RPL9 | Other |

**Table 7**

| **Genes Downregulated in Asymptomatic Females Compared to Asymptomatic Males** | |
| --- | --- |
| **Genes** | **Function** |
| CCL22 | Immune Response |
| SERPINE1 | Repair |
| SDC4 | Extracellular Matrix |
| COL9A3 | Extracellular Matrix |
| CPE | Neural |
| CYorf15A | Other |
| SNORD18C | Other |
| CYorf15B | Other |
| FOXQ1 | Other |
| HERC2P2 | Other |

**Table 8**
